# Functional analysis of caspase cleavable proteoforms from the *Drosophila* GSK-3 gene *shaggy*

**DOI:** 10.1101/2020.12.22.423955

**Authors:** Dagmara Korona, Daniel Nightingale, Bertrand Fabre, Michael Nelson, Bettina Fischer, Glynnis Johnson, Simon Hubbard, Kathryn Lilley, Steven Russell

## Abstract

The *Drosophila shaggy* (*sgg*) gene encodes the major fly orthologue of Glycogen Synthase Kinase −3 (GSK-3), a key highly conserved kinase at the heart of many signalling pathways. The *sgg* locus is complex, encoding multiple protein isoforms that are expressed in distinct temporal and tissue-specific patterns across development. Its isoforms predominantly differ at the carboxy and amino termini due to the use of different transcriptional start sites and alternative splicing events that include internal and terminal exons. One interesting class of proteins isoforms is represented by the Sgg-PD class (Sgg46), three proteoforms that contain a large 582 amino acid N-terminal domain which contains recognition sites for caspase-mediated cleavage. Regulated cleavage at these sites by non-apoptotic caspases has previously been implicated in the regulation of Sgg activity in adult bristle development. Here, we take a genome engineering approach to introduce specific tags into this unique Sgg-PD exon and utilise these for localisation and protein interaction studies. We also generated new loss of function alleles and specific mutations in the caspase cleavage motifs. We find that loss of functions Sgg-PD class alleles are viable and fertile, but exhibit adult locomotor and bristle defects. Expression analysis of lines carrying tags on both sides of the caspase cleavage sites indicates that the cleavage is developmentally regulated during embryogenesis. Surprisingly, we found that in some cells, particularly embryonic hemocytes, the N-terminal domain released by caspase cleavage is retained while the polypeptide containing the conserved kinase domain is apparently lost. Transcriptomic analysis of embryos homozygous for the new caspase-insensitive allele indicates a role for Sgg-PD in the regulation of cytoskeletal and cell junction functions, which is supported by proteomics analysis using specific in locus tags to identify common and unique protein interaction partners with N- and C-terminal domains. Taken together, our work identifies new activities for the Sgg protein and uncovers unexpected roles for caspase cleavage in Sgg biology.

## Introduction

The *Drosophila shaggy* (*sgg*) locus encodes the highly conserved protein kinase, Glycogen Synthase Kinase-3 (GSK-3), that is central to several signalling pathways in both vertebrates and invertebrates. Along with a key role as a negative regulator of the Wnt pathway, GSK-3 has been implicated in the Hedgehog, Insulin and G-protein-coupled receptor pathways, where it contributes to diverse developmental processes and is associated with multiple human diseases (1–5). Sgg contributes to the regulation of cell proliferation, apoptosis and morphogenesis via control of gene transcription, protein translation, cytoskeletal organization and the cell cycle (6,7). In *Drosophila*, the *sgg* locus is complex, producing at least 17 transcript variants that encode 10 different proteoforms (Figure 1A). We previously reported an analysis of *sgg* complexity by introducing isoform-specific tags into the endogenous locus, concluding that each proteoform has a unique spatial and temporal distribution across embryonic development (8). A functional analysis of two major proteoform classes, Sgg-PA and Sgg-PB, led us to conclude that these may represent the orthologues of the GSK-3α and GSK-3β proteins encoded by separate genes in vertebrates. We also reported the expression of a class of proteoforms previously described as Sgg46 (9), that are characterised by a 582 amino acid C-terminal domain unique to three proteins encoded by the locus (Sgg-PD, Sgg-PP and Sgg-PQ). The Sgg-PD class N-terminal domain contains two canonical caspase cleavage motifs (DEVD^235^ and DEVD^300^, Figure 1B) (10) but has no other recognisable protein domains within this region. Cleavage at these sites is predicted to release a 232 amino acid N-terminal polypeptide and a Sgg proteoform containing the conserved kinase domain with a 280 amino acid N terminal extension unique to this proteoform class. The entire Sgg-PD N-terminal domain is apparently unique to the Drosophilids, and although related sequences can be found in the *sgg* locus of other flies, including the house fly and stable fly, these lack the caspase cleavage sites. Related sequences have not been detected in the genomes of other invertebrates or vertebrates.

**Figure 1.**
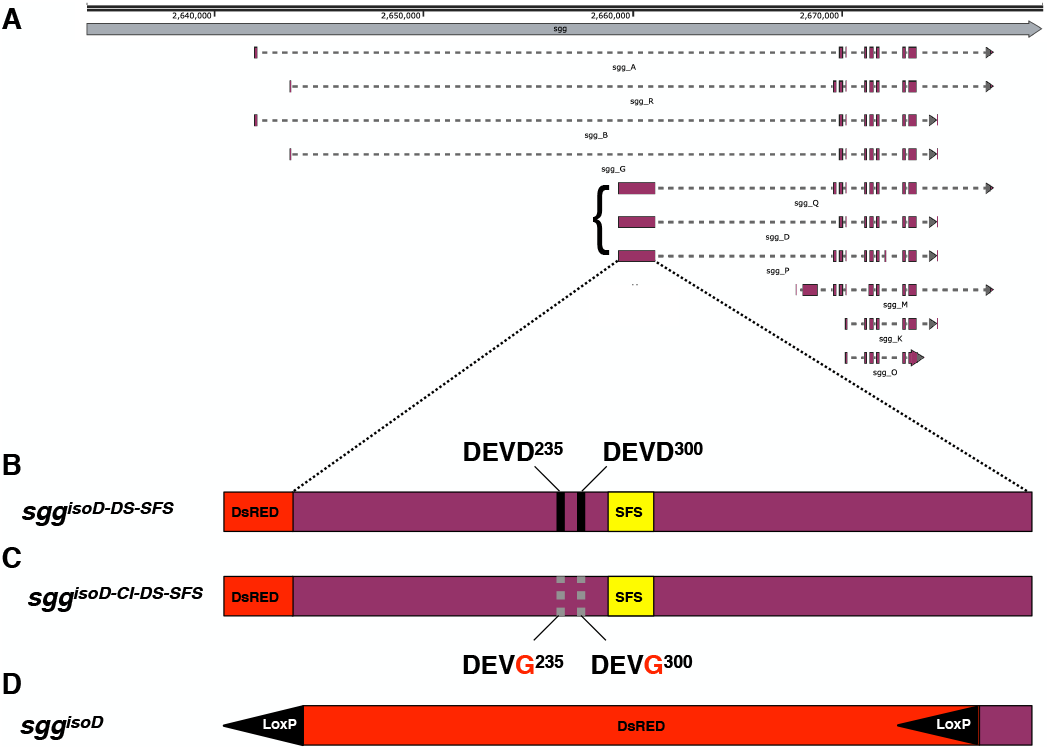
The sgg locus. **A**. Cartoon of the *sgg* locus on the *X* chromosome, numbers on the top line indicate nucleotide position. The structures of the 10 major Sgg proteoforms are indicated with the Sgg46 class highlighted with the bracket. **B**. The expand region represents the Sgg-PD class specific N-terminal exon (Sgg-PD, -PP and -PQ) with the location of the caspase cleavage sites indicated with black vertical lines. The locations of the DsRed and StrepII-Flag-StrepII tags in the engineered *sgg*^*isoDTag*^ line are indicated. **C**. The *sgg*^*isoD-CI*^ line with mutated caspase sites. **D**. In the *sgg*^*isoD*^ null allele the coding exon is replaced by the DsRed coding sequence driven by a 3XP3 promoter flanked by LoxP sites.

Analysis of mRNA expression by modENCODE (11) indicate that transcripts containing the exon are contributed maternally and expressed throughout development. We previously characterised the expression of proteoforms carrying this N-terminal domain using exon-specific protein tags introduced into the *sgg* locus via CRISPR-Cas9 genome engineering. We observed early ubiquitous expression that was subsequently elevated in the developing mesoderm, followed by the CNS, with prominent expression in the salivary glands, Malpighian tubules and proventriculus toward the end of embryonic development (8). Previous characterisation of the *sgg* locus indicated that the Sgg-PD class is not required for viability but can provide some functional rescue of null alleles (9). Subsequent work by Kanuka and colleagues (12) identified a role for Sgg-PD in sensory organ precursor formation. In particular, they found that a caspase-dependent cleavage of the Sgg-PD N-terminal exon was required to activate Sgg kinase activity, inhibit Wingless signalling and promote development of macrochaeta on the adult thorax. The analysis indicated that the uncleaved form was not active and concluded that developmental modulation of the caspase cleavage contributed to the function of Sgg-PD. It is therefore of some interest to determine any functional roles this protein class may have in *Drosophila*.

To help address the role of Sgg-PD class proteoforms and further investigate the contribution of the caspase cleavage, we adopted a CRISPR-Cas9 based genome engineering strategy to tag specific Sgg-PD proteoforms. Introducing multiple fluorescent protein or protein affinity tags into the N-terminal exon defining the three Sgg-PD class proteins allowed us to characterise the expression and protein interactions of both the polypeptides generated by caspase cleavage. We also generated new in locus mutations, either deleting the majority of the exon to produce a null allele for this proteoform class or specifically mutating the caspase cleavage sites. Using these mutant lines we confirm that the Sgg-PD class is not required for viability but has a role in aspects of nervous system function. Our analysis indicates that caspase cleavage is important in modulating Sgg-PD activity, confirming its role in adult sensory organ precursor development (12). We find expression and proteomic evidence that indicates the N-terminal domain generated by caspase cleavage is retained in some cells of the embryo and shows a unique repertoire of protein interactions. Our analysis connects Sgg-PD proteoforms to aspects of embryonic hemocyte biology, regulation of the cytoskeleton and homeostasis. Our work further emphasises the complexity of this key protein kinase and the diverse roles it plays in development.

## Results and Discussion

### Sgg-PD class alleles

In order to explore the role of the Sgg-PD family of proteoforms and how caspase cleavage of the 582 amino acid N-terminal domain unique to this proteoform class impacts any functions, we adopted a CRIPR-Cas9-based genome engineering approach to generate modifications at the endogenous *sgg* locus. First, we introduced tags either side of the two predicted caspase cleavage sites: at the N terminus we added a DsRed tag and downstream of the consensus caspase sites we added an in-frame StrepII-Flag-StrepII (SFS) tag (13). The tagged exon was introduced into the *sgg* locus by homology directed repair and we designate this allele *sgg*^*isoD-DS-SFS*^, which we will refer to as *sgg*^*isoDTag*^ for brevity. We generated a second repair template where the terminal aspartic-acid residues in two predicted DEVD caspase sites cleavage sites were mutated to glycines, rendering the exon insensitive to caspase activity, and designate this allele *sgg*^*isoD-CI-DS-SFS*^ which we refer to as *sgg*^*IsoD-CI*^. Finally, we created a null allele, *sgg*^*isoD*^, replacing the majority of the exon with a Ds-Red marker cassette (**Figure 1**). Although we observed very occasional segmentation defects in the null or caspase insensitive lines (<10%), overall the three lines we generated are homozygous or hemizygous viable and fertile.

Our previous analysis with an N-terminal SFS tagged version of Sgg-PD revealed expression of this proteoform family in the mesoderm from stage 9 of embryonic development, followed by elevated levels in the nervous system and prominent expression in the salivary glands (8). We analysed the expression of our new dual tag line by fluorescence microscopy imaging of briefly fixed embryos following the N terminal portion of the exon via DsRed fluorescence and the portion downstream of the caspase cleavage sites with Alexa488 labelled anti-Flag (**Figure 2**). We expected that in places where the exon is not subject to caspase cleavage, or is cleaved and both fragments are retained, we would observe both Ds-Red and Alex488 signals. If the exon is cleaved and the N-terminal portion degraded we expect only the Alexa488 signal and finally, if the N terminal peptide is retained after cleavage but the rest of the proteoform degraded we expect only a Ds-Red signal. As a control for the caspase cleavage we also examined the expression of both tags in the caspase insensitive variant.

**Figure 2.**
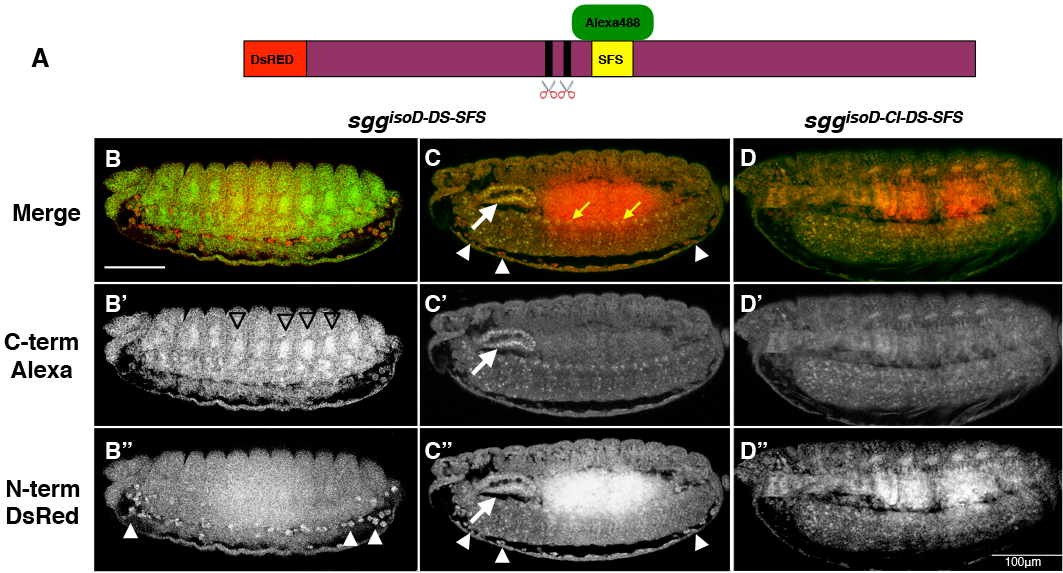
Expression of Sgg-PD proteoforms. **A**. Cartoon of the first exon of Sgg-PD class proteoforms with the location of N-terminal DsRed tag and the SVS tag downstream of the caspase cleavage sites (indicated by the scissors), the latter detected with Alexa488 - anti-FLAG. **B and C**. Expression of Sgg-PD proteoforms at stage 16 (B) and 14 (C). Triangles in B’ indicate developing mesoderm, triangles in B” indicate hemocytes. White arrow indicates the salivary gland, triangles indicate hemocytes and yellow arrow indicates cells in the CNS. **D**. Stage 15 embryo from the caspase insensitive line showing very similar expression of DsRed and Alexa488. The intense signal in the DsRed channel is due to gut autofluorescence. All embryos are lateral views with anterior to the left. Scale bar in B = 100 μm.

We found that the predominant signal across the embryo represented detection of both tags, indicative of no cleavage or cleavage events that resulted in the retention of both fragments, this is most clearly observed in the salivary gland where we previously reported high expression (**Figures 2B and C**). However, we also found examples of elevated levels of Alexa488 in the developing nervous system and mesoderm, indicative of cleavage and loss of the N-terminal fragment. Finally, we found highly specific Ds-Red only signal associated with what appears to be a subset of embryonic hemocytes (**Figure 2B”**), indicative of elevated levels of the N-terminal fragment post caspase cleavage with a loss of the remainder of the protein. The relatively uniform signals with Ds-Red and Alexa488 in the caspase insensitive *sgg*^*isoD-CI*^ line supports the view that the differential localisations we observe are due to caspase cleavage activity (**Figure 2D**). These observations confirm that endogenous Sgg proteoforms that include the caspase cleavable domain are subject to cleavage *in vivo*, that the cleavage shows tissue-specific regulation and, of particular interest, in some cells the N-terminal domain appears to be preferentially retained after caspase cleavage.

To confirm the expression in embryonic hemocytes, we examined the expression of Ds-Red in *sgg*^*isoDTag*^ and *sgg*^*IsoD-CI*^ lines along with antibody staining for the fascin hemocyte marker Singed (Sn) (14). As shown in **Figure 3**, the DsRed marker colocalises with Sn in hemocyte cells throughout the embryo, specifically the highly migratory plasmatocytes. We found that this co-localisation is not dependent on caspase cleavage, since expression in the caspase insensitive line appears to be identical to that observed in the cleavable wild type allele. Furthermore, the distribution of hemocytes appears unchanged in the caspase insensitive line, indicating that caspase cleavage is not necessary for the proper development of these cells. These observations further support the idea that caspase cleavage of Sgg-PD proteoforms is regulated in a tissue specific fashion (12). It remains to be determined, whether the hemocytes in the caspase insensitive line are biochemically or functionally normal, however, our expression profiling data described below suggest that the cleavage, and by inference the N-terminal domain lacking Sgg kinase activity, are involved in the normal activity of embryonic hemocytes.

**Figure 3.**
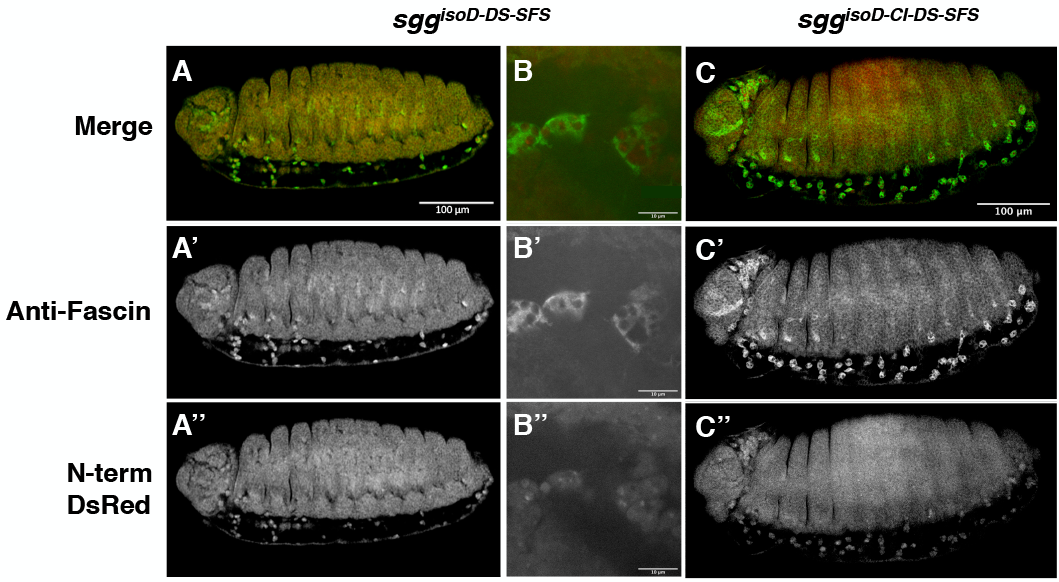
Expression of Sgg-D in embryonic hemocytes. **A**. Lateral view of stage 16 Sgg-PD tagged embryo imaged for DsRed (A”) and with anti-fascin (A’) showing co-localisation in hemocytes. **B**. Close up of hemocytes showing co-localisation. **C**. Stage 16 embryo carrying DsRed tagged caspase insensitive Sgg-PD variant imaged for DsRed and stained with anti-fascin showing continued co-localisation in hemocytes.

We next examined adults homozygous or hemizygous for the caspase insensitive *sgg*^*isoD-CI*^ allele and the *sgg*^*IsoD*^ null allele for any morphological phenotypes. Concordant with work employing ectopic expression of a caspase insensitive Sgg-PD variant (12), we observed the presence of extra anterior scutellar bristles in approximately 30% of adult flies (**Figure 4A-C**), compared to the 19% previously reported with the ectopic expression assay. We observed similar phenotypes with the null allele, supporting the previous conclusions of Kanuka and colleagues (12) that caspase cleavage was necessary to activate Sgg activity in the pathway. We also noticed defects in the adult arista, with approximately 13% of caspase insensitive mutants showing mild ectopic branching (**Figures 3D-F**). Interestingly, it has previously been shown that non-apoptotic caspase activity regulates arista morphology (15) and our observations suggest that, at least in part, this may operate through cleavage of Sgg-PD.

**Figure 4.**
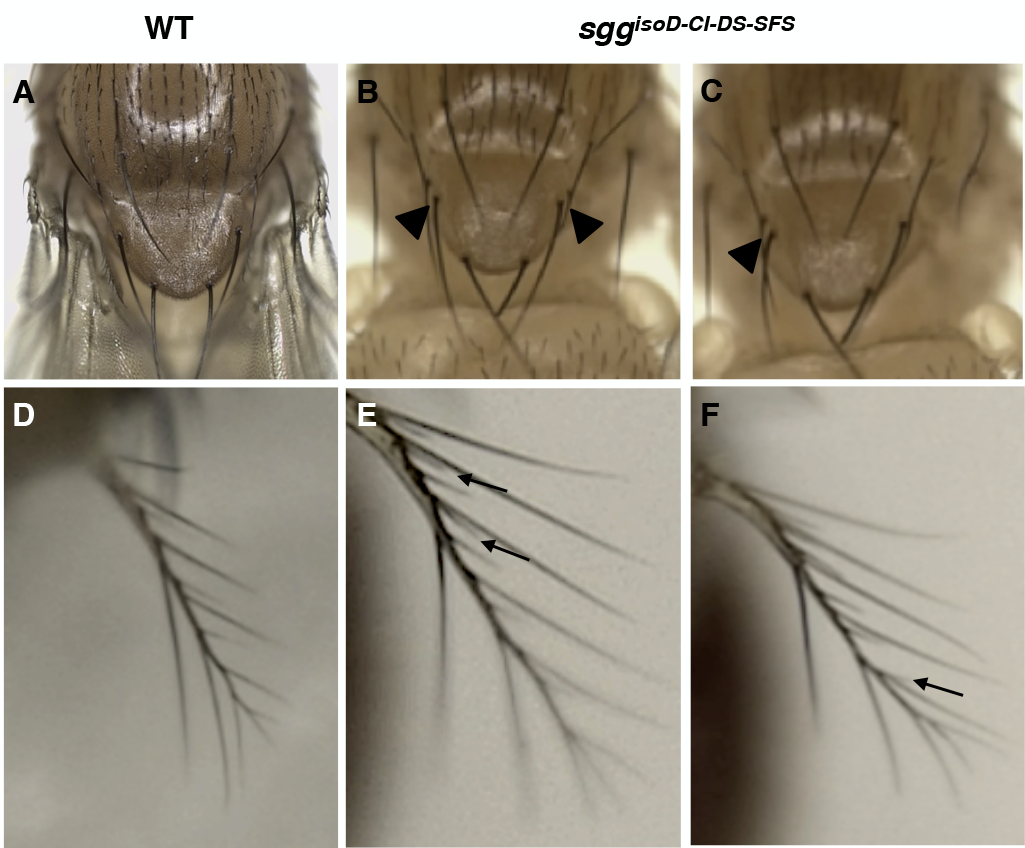
Sgg-PD caspase cleavage dependent phenotypes. **A**. Dorsal thorax and scutellum of the wild type. **B and C**. Dorsal thorax and scutellum of the caspase insensitive *sgg*^*isoD-CI*^ line showing duplicated anterior scutellar bristles (triangles). **D**. Wild type adult arista. **E and F**. Arista from the caspase insensitive *sgg*^*isoD-CI*^ line showing extra branching (arrows).

The role of GSK3 in the aging process has been the subject of some debate, with conflicting evidence emerging from different studies (16–18). In *Drosophila*, studies employing RNAi knockdown of *sgg* report shortened lifespan (18) and our recent work with *sgg*^*isoA*^ specific mutations supports this view (8). To identify any role for Sgg-PD proteoforms in longevity we performed standard adult survival assays with the *sgg*^*isoD*^ and *sgg*^*isoD-CI*^ mutants, comparing them with wild type flies from a matched genetic background (**Figure 5A)**. Homozygous females and hemizygous males were examined separately. With the *sgg*^*isoD*^ null allele we found significantly reduced survival (∼10%, p<0.05) in both sexes and for the caspase insensitive allele we found a similar lifespan reduction in males but a much weaker effect in females (∼6%, p <0.05). Our previous work with Sgg-PA specific mutants found a more sever lifespan reduction (15% in females, 25% in males), suggesting that different Sgg proteoforms contribute to adult longevity. Since Sgg-PD is expressed in the developing nervous system and our previous work identified adult locomotor defects with Sgg-PA specific mutants (8), we examined our *sgg* alleles in an adult climbing assay. We found strong reduction in locomotor activity of ∼35% in females and ∼26% in males at 10 days old (**Figure 5B)**, with a similar effect seen in the both null and caspase insensitive alleles. Taken together, these observations indicate that loss of the Sgg-PD class of proteoforms has defined developmental and adult homeostasis functions that in part are due to loss of the entire N-terminal domain and, to a lesser extent, rely on a caspase cleavage event. Our analysis of the embryonic expression suggests that in some circumstances these functions may be specifically mediated by the N-terminal domain after caspase cleavage. Unfortunately, the amino acid sequence of the Sgg-PD exon or the cleaved N-terminal domain provide no clues as to potential functions with no recognizable protein domains.

**Figure 5.**
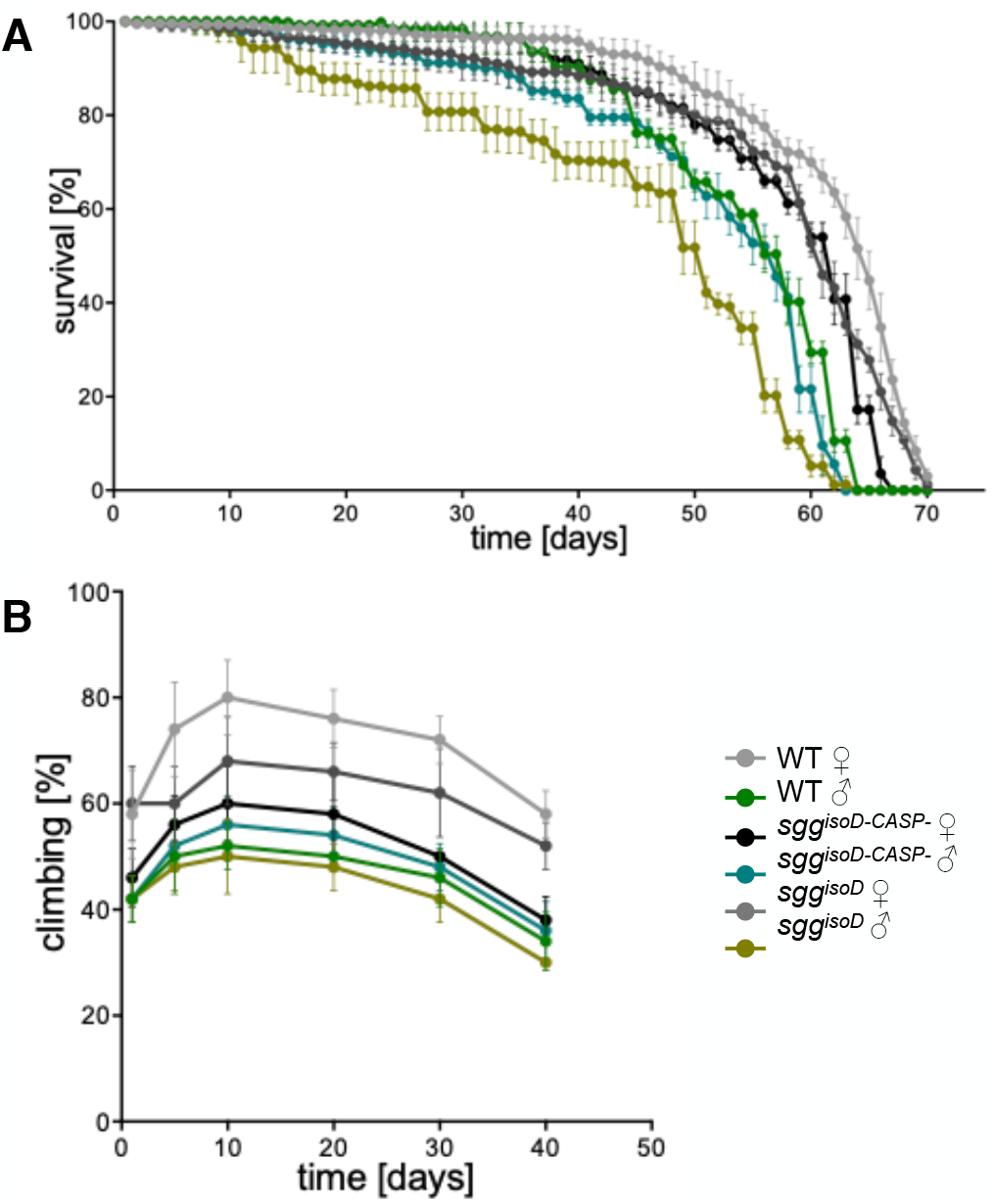
Sgg-PD adult phenotypes. **A**. Graph of lifespan from replicate lines of progenitor controls, *sgg*^*isoD*^ null and *sgg*^*isoD-CI*^ females and males as indicated in the key below. Error bars represent standard deviation. **B**. Graph of locomotor activity as measured by climbing assays with replicate lines of progenitor controls, *sgg*^*isoD*^ null and *sgg*^*isoD-CI*^ females and males as indicated. Error bars represent standard deviation.

### Transcriptome analysis

To gain insights into potential biological roles for the Sgg-PD caspase cleavage, we performed replicated RNA-seq analysis comparing RNA extracted from *sgg*^*isoD-Tag*^ embryos with stage matched *sgg*^*isoD-CI*^ caspase insensitive embryos. The analysis revealed 932 genes with significant expression changes (+/- 1.6-fold, adjusted p < 0.05) of which 737 were upregulated and 195 downregulated (Table S1). A gene ontology enrichment analysis identified approximately 70 genes associated with cuticle development that were downregulated in the mutant (adjusted p = 9.7E-33), most of which encode structural components, including 25 *Cpr* genes and 11 *Lcp* genes. An examination of the upregulated genes found a significant enrichment for genes involved in apical cell constriction, specifically a set of five Zona Pellucida Domain (ZPD) proteins involved in cuticle development (adjusted p = 6.9E-6) (19). In the embryo, these five genes (*m, mey, neo, nyo* and *tyn*) are regulated by *shavenbaby* (*svb*), which is itself negatively regulated by Wnt signalling, providing a link between loss of *sgg*^*isoD*^ functions and the changes in gene expression. At least 3 of these ZPD genes have phenotypes in macrochaetae, consistent with our observations on adult phenotypes. We found an addition 45 genes reported to be regulated by *svb* (20) that were also affected in the caspase insensitive Sgg-PD mutant embryos, further emphasising the link between this Sgg proteoform and the *svb* pathway.

To determine if the gene expression effects we observed are specific to *sgg*^*isoD-CI*^, we compared the results of our analysis with our previous RNA-seq study of *sgg*^*isoA*^ (viable) and *sgg*^*isoB*^ (lethal) mutant embryos (8), identifying 44 and 230 overlapping genes respectively. As we previously reported, there was no significant enrichment of any functional classes with the genes affected by loss of *sgg*^*isoA*^ and this is also the case for the 44 genes overlapping with *sgg*^*isoD-CI*^. In the case of *sgg*^*isoB*^, we found 47 downregulated cuticular proteins in common to the two alleles, all structural components of the cuticle. The overlap does not include the ZPD encoding genes highlighted above nor does the *sgg*^*isoB*^ gene list substantially overlap with the *svb* regulated gene list. Finally, we were interested in the cleavage dependent localisation of the N-terminal domain of Sgg-PD to hemocyte cells and compared our RNA-seq data with a list of genes reported to be expressed in embryonic hemocytes (21). We identified 195 genes misregulated in *sgg*^*isoD-CI*^ mutant embryos that show significant expression in hemocytes, including a set of structural cuticular proteins also regulated by *svb*. Taken together with our previous work on Sgg proteoforms, our RNA-seq analysis supports the view that individual proteoforms have unique functions. The identification of proteins involved in the embryonic epidermal cell biology controlled by the *svb* pathway specifically implicate the caspase cleaved Sgg-PD class of proteoforms in this process and provide a link with the cuticle phenotypes we observe in mutant adults. The observation that a significant number of genes misregulated in the caspase cleavage insensate mutants are expressed in embryonic hemocytes supports the view that the N-terminal domain of Sgg-PD has a role in hemocyte biology.

### Sgg-PD protein interactions

We next adopted a proteomics strategy to identify proteins interacting with specific Sgg-PD proteoforms. We used the iPAC approach with the specific tags introduced into the locus (22), performing replicated immunopurification followed by tandem mass-spectrometry analysis. We first used anti-DsRED, anti-FLAG and Streptactin to purify proteins from the *sgg*^*isoD-Tag*^ line to identify proteins interacting with Sgg-PD proteoforms. We reasoned that interactors identified with the N-terminal DsRED and the internal SFS tags should represent proteins associated with the uncleaved form, whereas interactors unique to each tag should represent proteins associated with the N or C terminal portions after cleavage. We performed a similar analysis using anti-DsRED to pull down interactors from the caspase insensitive *sgg*^*isoD-CI*^ line. As a control, we performed pull-downs with an untagged *w*^*1118*^ line. Using the QProt pipeline (23), applying an adjusted p-value cut-off of 0.05 and a requirement that interacting proteins were identified in at least 2 out of 3 independent pull downs after correction with wild type controls, we identified 79 proteins with the DsRED N-terminal bait and 102 with the SFS bait in the *sgg*^*isoD-Tag*^ line, and 188 with the DsRed pulldown from the *sgg*^*isoD-CI*^ line (**Figure 6)**.

**Figure 6.**
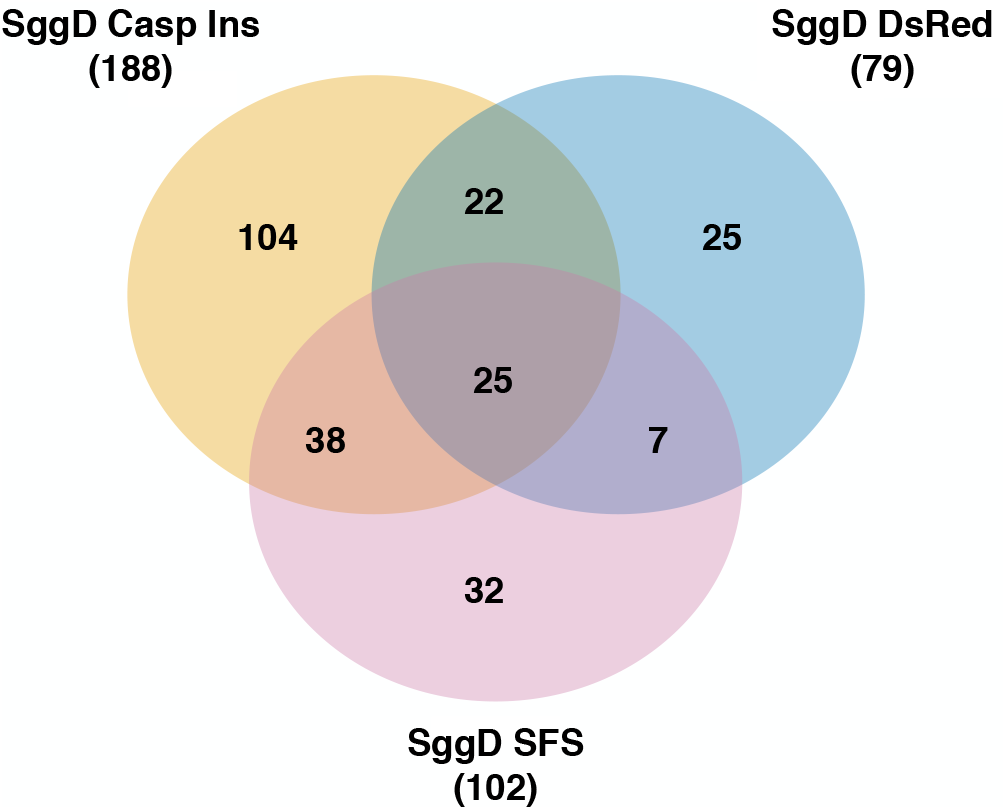
Overlap between proteomics studies. Venn diagram representing the overlap between the significant proteins identified in each of the three indicated pull-down experiments. Numbers in brackets refer to the total number of significant proteins identified in each replicated pull down.

Summarising the analysis, we identified 250 unique proteins across the three experiments, with pulldowns using the DsRED N-terminal and SFS baits in the *sgg*^*isoD-Tag*^ line, and DsRed pulldown from the *sgg*^*isoD-CI*^ line, 26 of which were common to the three. We found an enrichment for proteins involved in cytoskeleton, cell junction and epithelial cell biology, with 64 proteins annotated with one of these terms. This included Alpha-Cat, the cadherin Shg, Cindr and the Notch pathway component Akap200, common to all three pulldowns, placing the Sgg-PD proteoform at the heart of cellular morphogenesis processes. The proteomics analysis was broadly consistent with our RNA-seq experiments, where we identified a significant upregulation of Zona Pellucida components in the caspase insensitive mutants, suggesting that as well as physically associating with cell junction protein complexes, Sgg-PD may play an active role in regulating their composition.

Turning to the specific analyses, we identified 32 proteins in common to the *sgg*^*isoD-Tag*^ DsRED and SFS pulldowns, implying they may interact with the uncleaved protein. Although there were no significant Gene Ontology enrichments, we did identify proteins associated with adherens junctions (Alpha-Cat, Cindr and Shg), two of which are components of the Catenin complex (Alpha-Cat and Shg), along with a further 6 proteins with functional annotations related to the cytoskeleton (Akap200, Cindr, Corn, Hts, Tral, Zormin). Of these, Hts, Shg and Zormin have previously been identified as physically interacting with Sgg according to the BioGRID database (24). These common interactions are consistent with known roles for Sgg and further emphasise a role for the Sgg-PD proteoform class in aspects of epithelial cell biology we inferred from our RNA-Seq analysis described above.

Looking at the interactions we specifically identified with either DsRED or SFS pulldowns, we infer that proteins identified with SFS but not DsRed represent interactions specific to the kinase domain containing C-terminal portion of Sgg-PD after caspase cleavage. We found 32 proteins specific to the SFS pulldown that were not identified in any other of our experiments (Table 1). Significantly, we found 13 proteins annotated to have roles in the cytoskeleton or cell junction, this included Mey and Neo, which we identified as misregulated in our RNA-seq screen. This strongly implicates the cleaved form as an important contributor to aspects of cytoskeletal cell biology. From the overall list of 102 proteins identified in SFS pulldowns, we observed a significant enrichment (adjusted p <2E-3) for Gene Ontology biological process or cellular component annotations relating to cell junctions (9 proteins), anatomical structure development (45 proteins) and cellular development (34 proteins). Along with the cytoskeletal associated proteins shared with the DsRED pulldown, we identified a further 4 proteins with similar annotations (CG42748, Lac, L(2)gl and Vari) as well as 13 proteins with signalling pathway annotations, including 6 components of the Beta-Catenin Destruction Complex pathway (Arm, CtBP, Prosbeta6, Rpn7, Rpn12 and Rpt2), including Beta-Catenin itself (Arm). We also found a number of proteins with annotated nervous system phenotypes, including 6 with reported chaetae phenotypes (Arm, Bif, Gnf1, Msi, Noc, Rala and Shg), providing a potential link to the adult phenotypes we describe above. Finally, since the SFS pulldown should identify proteins that interact with the kinase domain common to all Sgg proteoforms, we compared the list of interactors with the proteins we previously identified as interacting with the Sgg-PA and -PB proteoforms (8). We found 24 proteins in common, including Arm, along with several of the cytoskeletal associated proteins described above. Taken together, we suggest that the identification of known signalling pathway components indicate the pull downs are likely to be reliable and further emphasise the role of the Sgg-PD proteoform in cytoskeletal and cell junction biology. Identification of proteins shared across multiple pulldowns in different engineered fly lines provides a high confidence set of *in vivo* Sgg interacting proteins that focus interest on aspects of cell polarity and cytoskeletal organisation.

**Table 1.**
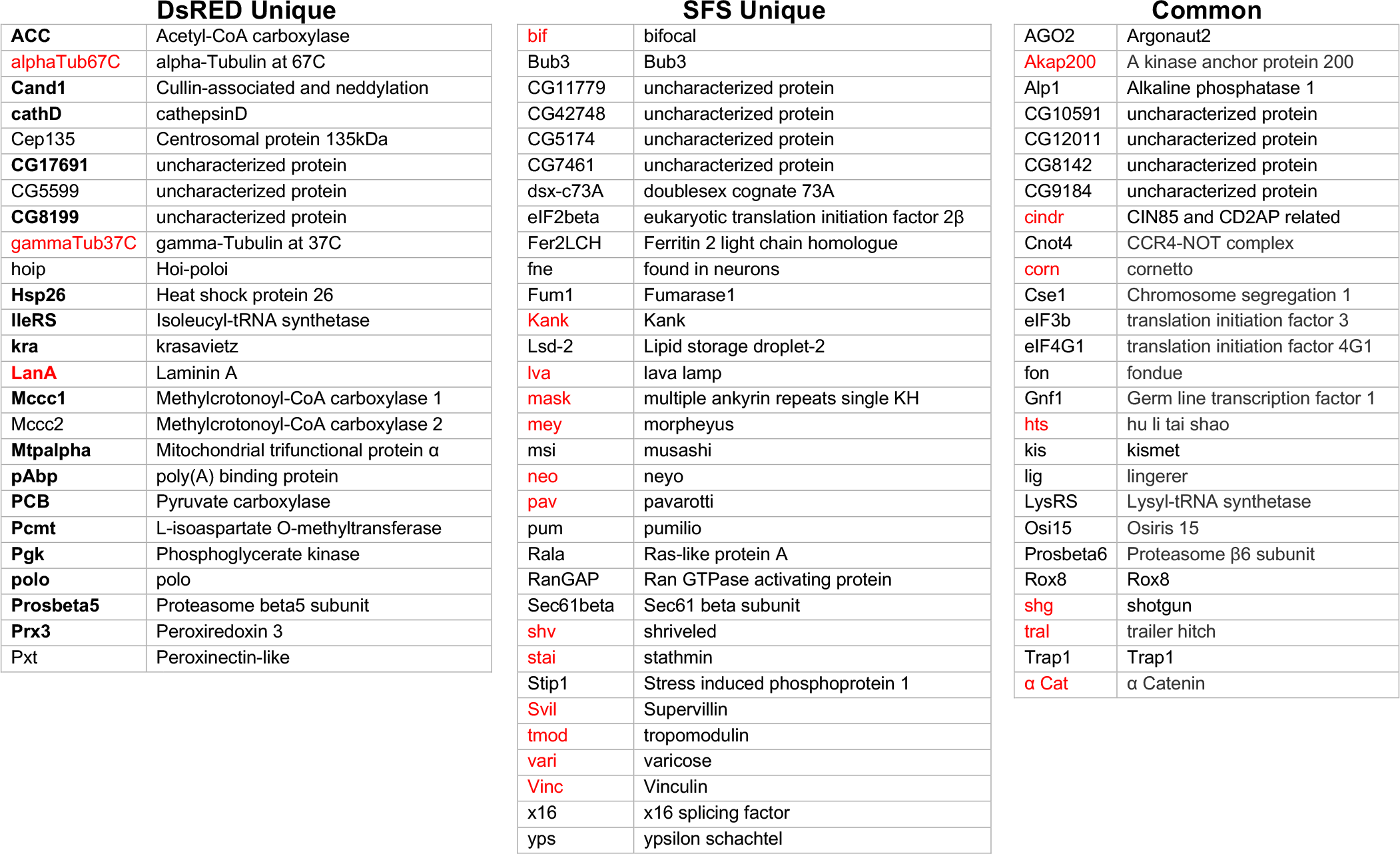
Sgg-PD interacting proteins. Gene symbols and names for proteins identified in Sgg-PD pulldowns. DsRED Unique = proteins specifically enriched by the N terminal DsRED tag that were not identified in other pull downs. SFS Unique = proteins specifically enriched by the StrepII-FLAG-StrepII tags downstream of the caspase cleavage sites that were not identified in other pull downs. Common = proteins identified in DsRED and SFS pulled downs from wild type, and DsRED pull downs from the caspase insensitive line. Red indicates annotations associated with cytoskeleton/cell junctions, bold indicates expressed in hemocytes.

Our imaging analysis indicated that the DsRed tagged N-terminal domain was maintained in some cells, particularly embryonic hemocytes, independent of the remainder of the proteoform after cleavage. Our proteomics identified 79 proteins with the DsRED pulldown, this list showed significant enrichment (p<8E-4) for proteins related to germ cell development (20 proteins), including some of the cytoskeletal components discussed above. However, we also found association with nuclear proteins (31) and ribonucleoprotein complexes (13). While our imaging suggests that the majority of the DsRED signal appears cytoplasmic, we cannot rule out some nuclear localisation. We note that this list includes proteasome components along with proteins known to have multiple cell locations, including cytoskeletal associations. It is therefore possible these interactions may represent involvement of Sgg-PD proteoforms in regulating intracellular transport. We found that 25 of the proteins were unique to the DsRED pulldown, 18 of which are expressed in embryonic hemocytes, consistent with our localisation data (Table 1), and 10 of the DsRed specific interactors are associated with carboxylic acid metabolism (p = 1.3E<5). We also note that both the DsRED-specific interactome and the set of genes expressed in embryonic hemocytes show a significant enrichment for targets of mir-277 (8 genes p=7.1E6 and 210 genes p=4.2E-5 respectively). The identification of DsRed specific interactors supports the idea that the N-terminal domain of Sgg-PD released after caspase cleavage is likely to have specific functions during embryonic development and in particular contribute to aspects of hemocyte biology.

Of the 188 potential interaction partners identified with anti-DsRED pull downs from the caspase insensitive *sgg*^*isoD-CI*^ line, 104 were not identified in our experiments with the wild type form. A total of 26 proteins were found in common between the caspase insensitive line and the two pulldowns from the wild type line, including the cell junction components α-Cat, Shg and Tra (Table 1). The proteins associated with caspase insensitive version showed significant enrichment for RNA metabolism pathways (47 proteins, p=1.8E-13) including rRNA processing enzymes, 5 translation initiation factors and a significant enrichment (p<1E-7) for proteins involved in mRNA stability. Whether these enrichments reflect defects in RNA processing and translation due to blocking the caspase-cleavage or a genuine enrichment of functional interactions that are stabilised by preventing cleavage remains to be determined. As well as pulling down Arm, the caspase insensitive version also identified the cell junction components (Lac, Alpha-Cat, l(2)gl, Cindr, Shg) we identified with the SFS pulldowns, strongly supporting the validity of these interactions since they were identified in a different fly line and using a different antibody for purification. This might also imply that these are bona fide interactions mediated by the C-terminal portion of Sgg-PD, distal to the caspase cleave site, since they are not identified with the N-terminal DsRED tag in the wild type: we would argue that this is also the case for the other 32 proteins shared between these two pull-down experiments. In terms of overlap between the anti-DsRED pull downs with the wild type and caspase insensitive lines, we found 22 shared proteins not identified by the SFS pulldown, including 12 of the proteins with nuclear annotations described above. Finding these in independent pulldown experiments strengthens our confidence in the interactions. We note with interest the association with Cse1, a protein involved in nuclear import that has a role in adult sensory organ development, consistent with the adult phenotypes we observe (15).

Our combined genetic, transcriptome and proteome analysis of new *sgg-PD* class alleles suggests a complex role for the Sgg-PD class of proteoforms in development and homeostasis, some of which appear to be regulated by caspase cleavage. A surprising finding, supported by imaging, RNA-seq and proteomics, is a role for the N-terminal 232 amino acid polypeptide released by caspase cleavage. We identified specific expression of this polypeptide in embryonic hemocytes with little or no retention of the C terminal kinase domain polypeptide.

## Materials and Methods

### Cloning gRNAs

To generate the transgenic fly lines carrying the tagged isoforms, we used CRISPR/Cas9 technology as previously described (25). We initially designed the insertion sites as indicated (Figure 1A) and choose appropriate gRNAs (Supp Table 3) that were cloned into pCDF3 (26). Briefly, target specific sequences were synthesized and either 5′-phosphorylated annealed, and ligated into the BbsI sites of pCDF3 pre-cut with BbsI. All gRNAs and primers are listed in Supp Table 3.

### Generation of donor vectors

For N-terminal tagging with DsRED and simultaneous internal tagging with 3xFLAG-StrepII in the Sgg-PD N-terminal domain containing caspase cleavage sites, a single donor vector was constructed to generate the *sgg-PD*^*isoD-DS-SFS*^ line. The inserted tags spanned the domain with the cleavage sites and included homology arms on both sites. To abolish DEVD caspase recognition sites site directed mutagenesis was performed at both sites: D235G (gat->ggt) and D300G (gat->ggt) using Gibson Assembly to generate a donor vector as described above to generate the *sgg-PD*^*isoDCI--DS-SFS*^ line. The *sgg*^*isoD*^ null allele was generated by replacing the first exon coding sequence with a cassette containing 3XP3 DsRed flanked with LoxP sites. Unless otherwise noted, cloning of donor vectors was performed with the Gibson Assembly Master Mix (New England Biolabs). PCR products were produced with the Q5 High-Fidelity 2X Master Mix (New England Biolabs). All inserts were verified by sequencing. Primers are listed in Supp Table 3.

### Drosophila methods

Embryos were injected using standard procedures into the *THattP40* line expressing *nos*-Cas9 (27,28). 500 ng/μl of donor DNA in sterile dH2O was injected together with 100 ng/μl of gRNA plasmid. Individually selected surviving adults were crossed to *w*^*1118*^ and the progeny screened for fluorescence: positively flies were balanced and homozygous stocks established for both caspase positive and negative variant. Injections were performed by the University of Cambridge Department of Genetics Fly Facility. All fly stocks were maintained at 25°C on standard cornmeal medium. Embryos were collected from small cages on yeasted grape juice agar plates.

### Locomotor behaviour

Adult female and male flies were collected shortly after eclosion and separated into 10 cohorts consisting of 10 flies (100 total) for each genotype. Flies were maintained at 25°C and transferred to fresh food every 3 days. For the climbing assay, each cohort was transferred to an empty glass cylinder (diameter, 2.5 cm; height, 20 cm), and allowed to acclimatize for 5 min. For each trial, flies were tapped down to the bottom of the vial, and the percentage of flies able to cross an 8-cm mark successfully within 10 s was recorded as the climbing index. Five trials were performed for each cohort, with a 1-min recovery period between each trial. Climbing assays were performed 1, 5, 10, 30 and 40 days after eclosion.

### Immunostaining and confocal microscopy

Localization of DsRED-tagged N-terminal fragment and 3xFLAG-StrepII tagged fragment following the caspase recognition sites of sgg were visualised in embryos. The former was visualized using native fluorescence, whereas the latter, was visualized by immunohistochemistry using Mouse Anti-FLAG M2 (F1804 Sigma), followed by Goat anti-Mouse IgG (H+L) Cross-Adsorbed Secondary Antibody, Alexa Fluor 488 (A-11001, Life Technologies) using standard protocols (29). Embryonic hemocytes were detected with an anti-Sn monoclonal from the Developmental Studies Hybridoma Bank (1:000 sn 7C) (30). Briefly, for fluorescence imaging, embryos were collected, dechorionated and quickly fixed to avoid bleaching of DsRED and stained with anti-FLAG or anti-Sn antibodies following standard protocols. Images were acquired using a Leica SP8 confocal microscope (Leica microsystems) with appropriate spectral windows for DsRED and Alexa488. Images were processed with the Fiji software (31).

### RNA seq

10-15hr embryos from a homozygous *sgg*^*isoD-CI-DS-SFS*^ stock and the tagged *sgg*^*isoD-DS-SFS*^ control line were collected and processed for RNA-seq as described below.

Tissue was homogenised in 300 μl TRIzol with a motorised pellet pestle for 30 seconds. The sample volume was then increased to 1 ml, then 200 μl of Chloroform was added and vortexed. Samples were centrifuged at max speed for 15 minutes at room temperature and the upper phase transferred to a new 1.5 ml tube. The RNA was then precipitated with 0.8 volumes of Isopropanol and incubated at −20°C for 2 hours. The samples were then centrifuged at 4°C at maximum speed for 30 minutes to pellet the RNA. The pellet was then washed with 1 ml 70% ethanol and centrifuged cold for another 5 minutes. After complete removal of the ethanol the RNA was dried for 5 minutes and re-suspended in 20 μl DEPC-treated water. The concentration of the samples was measured using the Qubit RNA HS Assay Kit and sample quality assess with the Bioanalyzer and or 1% agarose gel. Sequencing libraries were prepared using the NEBNext® Ultra™ II Directional RNA Library Prep Kit for Illumina. For each sample 100ng total RNA were processed with the NEBNext Poly(A) mRNA Magnetic Isolation Module. For all reactions, half reactions/volumes were used throughout the protocol, with exception for the AMPure bead clean-up step where washes were performed used the standard 200 μl 80% fresh Ethanol. Samples were barcoded and PCR amplification were performed for 12 cycles. After Bioanalyzer quality control equal amount of sample libraries were pooled and submitted for Illumina single-end sequencing.

Fastq reads were aligned using tophat (v2.1.1) with Bowtie (version: 2.3.4.0) using the default parameters. Gene counts tables across samples were created with Rsubread (1.22.3) using dmel_r6.20.gtf and options GTF.featureType=“exon” and GTF.attrType=“gene_id” and default parameters. Read were imported into edgeR (3.14.0) and filtered using the filterByExpr function with the default parameters. Data was normalised in limma (3.28.21) using limma-voom. Significant genes were identified fitting a linear model (lmFit) and empirical Bayes method (eBayes) (32–35). Genes were considered significant differential expressed with fdr <= 0.05 and logFC >= |0.7|. RNA-seq data are available from the NCBI Gene Expression Omnibus (GSE157625).

### iPAC-MS

Embryos from 8-20 hr collections were washed from agar plates with tap water, collected in 100 μm sieves, rinsed in the same solution to remove any yeast, dechorionated in 50% bleach for 1 min, rinsed again and placed on ice. Where necessary, washed embryos were frozen at −80°C until a sufficient quantity was collected. For each purification, ∼200 μl wet-volume of embryos were manually homogenized with a 2 ml Dounce homogenizer in 1 ml of extraction buffer (50 mM Tris, pH 7.5, 125 mM NaCl, 1.5 mM MgCl2, 1 mM EDTA, 5% Glycerol, 0.4% Igepal CA-630, 0.5% digitonin and 0.1% Tween 20) and processed essentially as previously described [13]. Samples were independently immunopurified using StrepII, FLAG and DsRED.

ANTI-FLAG^®^ M2 affinity gel (Sigma) and ***Strep***-Tactin® Superflow® resin (IBA) were used to capture each FLAG-tagged and StrepII-tagged bait and its binding partners, respectively (22). For pulldown of fluorescently tagged proteins (DsRED), anti-RFP mAb agarose resin (MBL International) was used. Briefly, protein concentration estimation in the embryo lysate was performed using a DC assay (Bio-Rad). The lysate was divided equally into three parts (6 mg total protein per pulldown), to which each resin, pre-washed in extraction buffer, was added. Following 2 h of incubation at 4°C on a rotating wheel, the resin was washed three times in extraction buffer. Immunoprecipitates were eluted twice each, using 100 μg/ml 3xFLAG peptide (Sigma) in lysis buffer for FLAG immunoprecipitates and 10 mM desthiobiotin in lysis buffer for *Strep*-Tactin immunoprecipitates; each for 10 minutes at 4°C. Anti-GFP resin immunoprecipitates were eluted in 100 mM glycine-HCl, pH 2.5 with gentle agitation for 30 seconds, followed by immediate neutralization in 1 M Tris-HCl, pH 10.4.

Purification of the baits was confirmed via immunoblots and samples were prepared for mass spectrometric analysis using in-gel digestion, allowing the sample to enter 2 cm into an SDS-PAGE gel. Gels were fixed and stained with colloidal Coomassie stain, after which the protein-containing band was excised and cut into two equally sized parts. Each band was destained, reduced with dithiothreitol, alkylated with iodoacetamide and subjected to tryptic digest for 16 hours at 37°C. Approximately 1 μg of peptides from each digested band was analysed using LC-MS/MS on a Q Exactive mass spectrometer (ThermoFisher Scientific), as previously described (36).

For label-free quantification (LFQ), data were processed using MaxQuant (version 1.6.3.4) (37). Raw data were searched against protein sequences contained in the all translation database obtained from FlyBase release FB2017_06 dmel_r6.19 at (ftp://ftp.flybase.net/releases/FB2017_06/dmel_r6.19/fasta/) (38). The database was customised with the addition of the Uniprot proteome for *Wolbachia pipientis wMel* (https://www.uniprot.org/uniprot/?query=proteome:UP000008215). Within MaxQuant, searches were performed against a reversed decoy dataset and a set of known contaminants. Default search parameters were used with digestion using trypsin allowing two missed cleavages and minimum peptide lengths of six. Carbamidomethyl Cysteine was specified as a fixed modification. Oxidation of Methionine, N-term protein Acetylation and Phosphorylation of Serine, Threonine and Tyrosine were specified as variable modifications. Additionally, “match between runs” was enabled with fractions specified to limit matching to occur only between replicates of the same bait and tag combination. Identification of significant interacting proteins was performed with the QProt pipeline (23). Gene Ontology enrichments were obtained from FlyMine (39) using the Benjamini-Hochberg multiple testing correction.

## Supplementary Material

***Supplementary Table 1. RNA-seq analysis***.

Table of genes showing significant expression changes comparing RNA from *sgg*^*isoDTag*^ embryos with stage matched *sgg*^*isoD-CI*^ caspase insensitive embryos. FBgn = FlyBase database identifier; Symbol = FlyBase gene symbol; Gene = FlyBase gene name; Log2 FC = differential expression value (Log2); Adjp = p value adjusted for multiple testing.

***Supplementary Table 2. Proteomics data***.

Table of genes encoding proteins significantly enriched in the indicated mass spectrometry analyses. FBgn = FBgn = FlyBase database identifier; Symbol = FlyBase gene symbol; Gene = FlyBase gene name. Sgg-PD DsRed ALL = all proteins enriched by the N-terminal DsRed pulldown from *sgg*^*isoDTag*^ embryos; Sgg-PD DsRed Unique = proteins only identified in the N-terminal pulldown from *sgg*^*isoDTag*^ embryos; Sgg-PD SFS ALL = all proteins enriched by the C-terminal Streptactin and FLAG pulldowns from *sgg*^*isoDTag*^ embryos; Sgg-PD SFS Unique = proteins only identified in the C-terminal pulldown from *sgg*^*isoDTag*^ embryos. Sgg-PD-CaspIns DsRED ALL = all proteins enriched by the N-terminal DsRed pulldown from caspase insensitive *sgg*^*isoD-C*^ embryos; Sgg-PD-CaspIns DsRED Unique = proteins unique to the DsRed pulldowns from *sgg*^*isoD-C*^ embryos.

***Supplementary Table 3. Primer and gRNA sequences used for vector construction.***

## Author Contributions and Notes

DK, DN, BF, MN, BF and SR performed experimental or analytical work. SR, SH and KSL conceived the study and raised the funding. DK and SR wrote the manuscript with all authors contributing. The authors declare no conflict of interest.

## Acknowledgments

We are indebted to Andrew Davidson and Will Wood, University of Edinburgh for reagents and invaluable insights into the world of hemocytes. We thank Boguslawa Korona, Dept of Pathology, University of Cambridge for preliminary Western blots. This work was supported by a BBSRC grant (BB/L002817/1) to SH, KL and SR. This document was formatted using a template from Ilya Finkelstein, University of Texas at Austin (https://github.com/finkelsteinlab/BioRxiv-Template).

## References

1. Bourouis M, Moore P, Ruel L, Grau Y, Heitzler P, Simpson P. An early embryonic product of the gene shaggy encodes a serine/threonine protein kinase related to the CDC28/cdc2+ subfamily. EMBO J. 1990; 9:2877–84.

2. Siegfried E, Perkins LA, Capaci TM, Perrimon N. Putative protein kinase product of the Drosophila segment-polarity gene zeste-white3. Nature. 1990; 345:825–9.

3. He X, Saint-Jeannet JP, Woodgett JR, Varmus HE, Dawid IB. Glycogen synthase kinase-3 and dorsoventral patterning in Xenopus embryos. Nature. 1995; 374:617–22.

4. Ali A, Hoeflich KP, Woodgett JR. Glycogen synthase kinase-3: properties, functions, and regulation. Chem Rev. 2001; 101:2527–40.

5. Kaidanovich-Beilin O, Woodgett JR. GSK-3: Functional Insights from Cell Biology and Animal Models. Front Mol Neurosci. 2011; 4:40.

6. Beurel E, Grieco SF, Jope RS. Glycogen synthase kinase-3 (GSK3): regulation, actions, and diseases. Pharmacol Ther. 2015; 148:114–31.

7. Patel P, Woodgett JR. Glycogen Synthase Kinase 3:A Kinase for All Pathways? Curr Top Dev Biol. 2017; 123:277–302.

8. Korona D, Nightingale D, Fabre B, Nelson M, Fischer B, Johnson G, et al. Characterisation of protein isoforms encoded by the Drosophila Glycogen Synthase Kinase 3 gene shaggy. PLoS One. 2020;15:e0236679.

9. Ruel L, Pantesco V, Lutz Y, Simpson P, Bourouis M. Functional significance of a family of protein kinases encoded at the shaggy locus in Drosophila. EMBO J. 1993; 12:1657–69.

10. McStay GP, Salvesen GS, Green DR. Overlapping cleavage motif selectivity of caspases: implications for analysis of apoptotic pathways. Cell Death Differ. 2008; 15:322–31.

11. Graveley BR, Brooks AN, Carlson JW, Duff MO, Landolin JM, Yang L, et al. The developmental transcriptome of Drosophila melanogaster. Nature. 2011; 471:473–9.

12. Kanuka H, Kuranaga E, Takemoto K, Hiratou T, Okano H, Miura M. Drosophila caspase transduces Shaggy/GSK-3beta kinase activity in neural precursor development. EMBO J. 2005; 24:3793–806.

13. Lowe N, Rees JS, Roote J, Ryder E, Armean IM, Johnson G, et al. Analysis of the expression patterns, subcellular localisations and interaction partners of Drosophila proteins using a pigP protein trap library. Development. 2014; 141:3994–4005.

14. Zanet J, Jayo A, Plaza S, Millard T, Parsons M, Stramer B. Fascin promotes filopodia formation independent of its role in actin bundling. J Cell Biol. 2012; 197:477–86.

15. Cullen K, McCall K. Role of programmed cell death in patterning the Drosophila antennal arista. Dev Biol. 2004; 275:82–92.

16. Castillo-Quan JI, Li L, Kinghorn KJ, Ivanov DK, Tain LS, Slack C, et al. Lithium Promotes Longevity through GSK3/NRF2-Dependent Hormesis. Cell Rep. 2016; 15:638–50.

17. Souder DC, Anderson RM. An expanding GSK3 network: implications for aging research. Geroscience. 2019; 41:369–82.

18. Trostnikov MV, Roshina NV, Boldyrev SV, Veselkina ER, Zhuikov AA, Krementsova AV, et al. Disordered Expression of shaggy, the Drosophila Gene Encoding a Serine-Threonine Protein Kinase GSK3, Affects the Lifespan in a Transcript-, Stage-, and Tissue-Specific Manner. Int J Mol Sci. 2019; 20:2200.

19. Fernandes I, Chanut-Delalande H, Ferrer P, Latapie Y, Waltzer L, Affolter M, et al. Zona pellucida domain proteins remodel the apical compartment for localized cell shape changes. Dev Cell. 2010; 18:64–76.

20. Menoret D, Santolini M, Fernandes I, Spokony R, Zanet J, Gonzalez I, et al. Genome-wide analyses of Shavenbaby target genes reveals distinct features of enhancer organization. Genome Biol. 2013; 14:R86.

21. Cattenoz PB, Sakr R, Pavlidaki A, Delaporte C, Riba A, Molina N, et al. Temporal specificity and heterogeneity of Drosophila immune cells. EMBO J. 2020; 39:e104486.

22. Rees JS, Lowe N, Armean IM, Roote J, Johnson G, Drummond E, et al. In Vivo Analysis of Proteomes and Interactomes Using Parallel Affinity Capture (iPAC) Coupled to Mass Spectrometry. Mol Cell Proteomics. 2011; 10:pM110 002386.

23. Choi H, Kim S, Fermin D, Tsou CC, Nesvizhskii AI. QPROT: Statistical method for testing differential expression using protein-level intensity data in label-free quantitative proteomics. J Proteomics. 2015; 129:121–6.

24. Oughtred R, Stark C, Breitkreutz BJ, Rust J, Boucher L, Chang C, et al. The BioGRID interaction database: 2019 update. Nucleic Acids Res. 2019; 47:D529–41.

25. Korona D, Koestler SA, Russell S. Engineering the Drosophila Genome for Developmental Biology. J Dev Biol. 2017; 5:16.

26. Port F, Chen HM, Lee T, Bullock SL. Optimized CRISPR/Cas tools for efficient germline and somatic genome engineering in Drosophila. Proc Natl Acad Sci USA. 2014; 111:E2967–76.

27. Ren X, Sun J, Housden BE, Hu Y, Roesel C, Lin S, et al. Optimized gene editing technology for Drosophila melanogaster using germ line-specific Cas9. Proc Natl Acad Sci USA. 2013; 110:19012–7.

28. Gratz SJ, Harrison MM, Wildonger J, O’Connor-Giles KM. Precise Genome Editing of Drosophila with CRISPR RNA-Guided Cas9. Methods Mol Biol. 2015; 1311:335–48.

29. Patel NH. Imaginal neuronal subsets and other cell types in whole mount Drosophila embryos and and larvae using antibody probes. Methods Cell Biol. 1994; 44:445–87

30. Cant K, Knowles BA, Mooseker MS, Cooley L. Drosophila singed, a fascin homolog, is required for actin bundle formation during oogenesis and bristle extension. J Cell Biol. 1994; 125:369–80.

31. Schindelin J, Arganda-Carreras I, Frise E, Kaynig V, Longair M, Pietzsch T, et al. Fiji: an open-source platform for biological-image analysis. Nat Methods. 2012; 9:676–82.

32. Trapnell C, Pachter L, Salzberg SL. TopHat: discovering splice junctions with RNA-Seq. Bioinformatics. 2009; 25:1105–11.

33. Liao Y, Smyth GK, Shi W. The R package Rsubread is easier, faster, cheaper and better for alignment and quantification of RNA sequencing reads. Nucleic Acids Res. 2019; 47:e47.

34. Robinson MD, McCarthy DJ, Smyth GK. edgeR: a Bioconductor package for differential expression analysis of digital gene expression data. Bioinformatics. 2010; 26:139–40.

35. Ritchie ME, Phipson B, Wu D, Hu Y, Law CW, Shi W, et al. limma powers differential expression analyses for RNA-sequencing and microarray studies. Nucleic Acids Res. 2015; 43:e47.

36. Dikicioglu D, Nightingale DJH, Wood V, Lilley KS, Oliver SG. Transcriptional regulation of the genes involved in protein metabolism and processing in Saccharomyces cerevisiae. FEMS Yeast Res. 2019; 19:foz014

37. Tyanova S, Temu T, Cox J. The MaxQuant computational platform for mass spectrometry-based shotgun proteomics. Nat Protoc. 2016; 11:2301–19.

38. Thurmond J, Goodman JL, Strelets VB, Attrill H, Gramates LS, Marygold SJ, et al. FlyBase 2.0: the next generation. Nucleic Acids Res. 2019; 47:D759–65.

39. Lyne R, Smith R, Rutherford K, Wakeling M, Varley A, Guillier F, et al. FlyMine: an integrated database for Drosophila and Anopheles genomics. Genome Biology. 2007; 8:R129.

